# Network analysis uncovers associations in the turnover of C1 molecules in a winter lake

**DOI:** 10.1101/2022.10.31.514531

**Authors:** Rhiannon Mondav, Gaëtan Martin, Sari Peura, Sarahi L Garcia

## Abstract

The generation and consumption of single carbon molecules (CO_2_ and CH_4_) by aquatic microbial communities is an essential aspect of the global carbon budget. Organic carbon flow (warm sunlit regimes) is depicted as beginning at the surface with autochthonous fixation followed by biomass settling to sediments, CO2 respiration to the atmosphere, and outflow. We sought to broaden understanding of C1 cycling and consortia by examining the microbial community of a below-ice lake water column in which both input and output are likely disrupted due to ice cover. By analysing the microbial community composition and co-occurrence network of an ice-covered lake timeseries, we were able to identify potential consortia involved in C1 cycling. The network confirmed known associations supporting the efficacy of such analyses but also pointed to previously unknown potential associations. Further and contrary to typical organic carbon flow under warm sunlit regimes, we found support for upward flow of recently fixed carbon in cold low-light conditions under-ice in winter.

## Introduction

Historically, autochthonous carbon flow in lakes is visualised as flowing from the water surface down, through a microbial loop, and then outwards via sedimentation, gas flux, or water outflow (Fig 1) (Tranvik *et al*., 2009; Martinez-Cruz *et al*., 2015; Limpert *et al*., 2020). In this view, autochthonous carbon is fixed by photoautotrophs (cyanobacteria, diatoms, macro-algae) occurring in surface waters. The fixed carbon is then cycled through the microbial loop (grey circle Fig 1) in which some will be released back to the atmosphere as CO2 (black arrow up Fig 1). while some settles through the water column to the sediments (green arrow down Fig 1). Microbes in the sediments can remineralise organic carbon thus releasing CO_2_ back into the loop, while methanogens can convert a variety of different carbon stocks to CH_4_ (red arrows up Fig 1). CH_4_ can then be used by water-column microbes or may escape as flux to the atmosphere. Carbon flux to the atmosphere consists mostly of single carbon (C1) molecules CO_2_ and CH_4_, both of which are greenhouse gasses.

**Figure 1.**
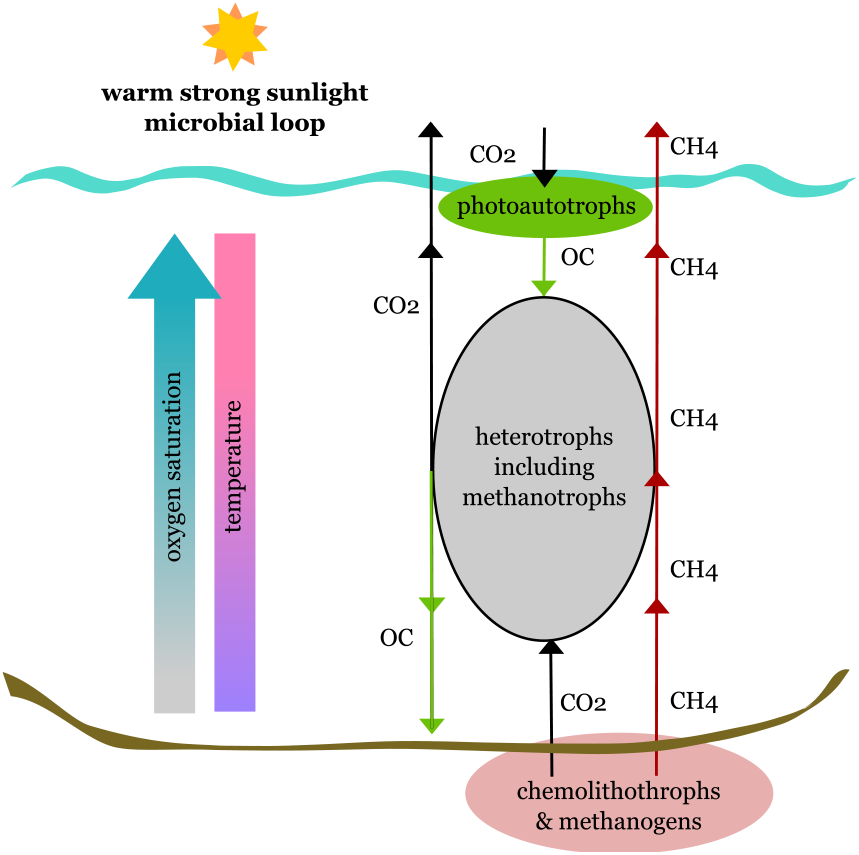
Diagram of standard perspective of lake microbial carbon loop

The ratio, activity, and location of lake microbes producing and consuming C1 molecules determine the flux of C1 greenhouse gases from lakes. For example, methane consuming microbes, methanotrophs, in the water column can oxidise up to 99 % of the CH_4_ produced in sediments before it reaches the atmosphere (Frenzel *et al*., 1990; Kankaala *et al*., 2006; Bastviken *et al*., 2011; Mayr *et al*., 2020). Alternatively, CH4 produced in the sediment can elude methanotrophs in the water-column by bubbling (ebullition) or transportation through plant tissue (Sebacher *et al*., 1985; Bastviken *et al*., 2008) thus reaching the atmosphere directly.

CH_4_ flux from inland waters is estimated to be 0.159 +/- 0.04 Pg/yr (Saunois *et al*., 2020) or around 3.9 Pg/yr CO_2_ 100 yr equivalents. CO_2_ flux from inland waters is estimated at somewhere between 2.1 +/- 0.4 Pg/yr (Raymond *et al*., 2013) and 58 Pg/yr (Pollard, 2022). Thus, the lowest estimated total flux of 6 Pg/yr CO_2_ 100 yr equivalents, is at least half that released from burning fossil fuels (Pollard, 2022). While lakes make up only one component of inland waters it is clear that understanding lake microbial C1 cycling is more important than their global surface area suggests. Because of the large flux of greenhouse gases between lakes and the atmosphere, it is important to have accurate up-to-date descriptions of carbon cycling in lakes.

Linking carbon cycling to lake microbial taxa is an important aspect of this description (Table 1). Assigning function based on taxonomy always suffers from problems of false positives (inaccurate taxonomic profile and presumptive functionality) and false negatives (incomplete metabolic profile). Table 1 presents an overview of known C1 cycling phyla, but, apart from Cyanobacteria and the rTCA cycle, other microbes should be classified to genus level before assigning putative C1 cycling capacity (SI Table 1, DOI: 10.6084/m9.figshare.21388005). Further, the detection of C1 cycling microbes in an environment might not correlate with activity and C1 flux is determined by the net ratio of fixation, oxidation, and reduction (Sawakuchi *et al*., 2021). Nevertheless, important insights into the lake C1 cycling can be made by assessing the location and abundance of putative C1 cycling microbes.

**Table 1.** Summary of paths to produce or consume CO2 and CH4, and affiliated phyla

| **C<sub>1(gas)</sub> cycle component** | **Pathway or enzyme** | **Taxa - phyla** | **References** |
| --- | --- | --- | --- |
| C fixation | reductive pentose phosphate cycle (Calvin-Benson-Bassham cycle) | *Cyanobacteria, Plantae, dinoflagellates, Bathyarchaeota, Pseudomonadota (Proteobacteria), Chloroflexota, Pseudomonadota, Actinomycetota* | (Levicán *et al.*, 2008; Grostern and Alvarez-Cohen, 2013; Alfreider *et al.*, 2017, 2018; Gaisin *et al.*, 2019; Pan *et al.*, 2020; Fang *et al.*, 2022; Taubert *et al.*, 2022) |
|  | reductive tricarboxylic acid cycle (rTCA) (Krebs) | *Nitrospirota, Chlorobi, Pseudomonadota, Campylobacterota, Deinococcota, Aquificaeota, Pseudomonadota, Bacillota* | (Kolinko *et al.*, 2016; Alfreider *et al.*, 2017, 2018; Cabello-Yeves *et al.*, 2018; Fang *et al.*, 2022) |
|  | reductive acetyl-CoA cycle (Wood-Ljungdahl) | *Euryarchaeota, Bathyarchaeota, Thorarchaeota, Lokiarchaeota, Heimdallarchaeota, Odinarchaeota, Aerophobetes, Desulfobacterota, Pseudomonadota, Bacillota* | (Länge *et al.*, 1988; Berg, Kockelkorn, *et al.*, 2010; Wang *et al.*, 2016; Huang *et al.*, 2019; Macleod *et al.*, 2019; Pan *et al.*, 2020; Usman *et al.*, 2021; Fang *et al.*, 2022) |
|  | 3-hydroxypropionate bicycle (3-HP) | *Chloroflexota* | (Herter *et al.*, 2001; Zheng *et al.*, 2022) |
|  | 3-hydroxypropionate/4-hydroxybutyrate pathway (HP/HB) | *Crenarchaeota, Thermoproteota, Thaumarchaeota, Euryarchaeota* | (Berg *et al.*, 2007; Berg, Kockelkorn, *et al.*, 2010; Berg, Ramos-Vera, *et al.*, 2010; Alfreider *et al.*, 2017, 2018; Abby *et al.*, 2018) |
|  | dicarboxylate/4-hydroxybutyrate cycle (DC/HB) | *Crenarchaeota, Thermoproteota* | (Huber *et al.*, 2008; Berg, Kockelkorn, *et al.*, 2010; Berg, Ramos-Vera, *et al.*, 2010) |
|  | reductive hexulose-phosphate cycle (RHP) | *Euryarchaeota* | (Kono *et al.*, 2017; Yang *et al.*, 2019; Zhang *et al.*, 2020) |
|  | reductive glycine pathway (rGly) | *Pseudomonadota* | (Figuerola *et al.*, 2018; Sánchez-Andrea *et al.*, 2020) |
| Methanogenesis | varied but all involve methyl-coenzyme M reductase (Mcr) | *Euryarchaeota*, *Bathyarchaeota*?, *Verstraetearchaeota*? | (Evans *et al.*, 2015; Wen *et al.*, 2017; Bräuer *et al.*, 2020) |
| Methane production | FeFe or VFe type nitrogenases | *Pseudomonadota*, *Bacillota*, *Euryarchaeota*, *Cyanobacteria*, *Bacteroidota*, *Chlorobiota*, *Firmicutes*, *Verrucomicrobiota* | (Masukawa *et al.*, 2009; McGlynn *et al.*, 2012; Mus *et al.*, 2018) |
|  | demethylation by C-P lyase, etc | *Actinomycetota*, *Pseudomonadota*, *Cyanobacteria*, *Chloroflexota* | (Dyhrman *et al.*, 2006; Martínez *et al.*, 2013; Carini *et al.*, 2014; Yao *et al.*, 2016; Bižić *et al.*, 2020) |
| Methane release | Fenton-driven reactions | *fast growing bacteria*, *Bacillota*, *Pseudomonadota*, *Euryarchaeota*, *Ascomycota*, *Plantae*, *Chordata* | (Ernst *et al.*, 2022) |
| Methane oxidation | involve Mcr and an alternative electron acceptor | *Euryarchaeota* | (Yu *et al.*, 2021; Chadwick *et al.*, 2022) |
|  | membrane-bound Cu (particulate) methane monooxygenase | *Pseudomonadota*, *Bacillota*, *Euryarchaeota*, *Cyanobacteria*, *Bacteroidota*, *Chlorobiota*, *Firmicutes*, *Verrucomicrobiota*, *NC10* | (Guerrero-Cruz *et al.*, 2021; Schmitz *et al.*, 2021) |
|  | cytosolic Fe (soluble) methane monooxygenase | *Pseudomonadota* | (Hakobyan and Liesack, 2020; Haque *et al.*, 2020) |
| CO<sub>2</sub> release | tricarboxylic acid cycle | *many* |  |
|  | pentose phosphate cycle | *many* |  |
|  | Over 1100 different reactions | *many* | (Caspi *et al.*, 2020) |

A factor that affects C1 cycling is the composition of the microbial community. More specifically cross-feeding and syntenic interactions between microbes can alter not just activity but presence by expanding habitat ranges. For example, obligately aerobic methanotrophs can be active in anoxic waters if they co-occur with photosynthetic organisms (Oswald *et al*., 2015) that release oxygen. But, discovering and pinpointing more such symbioses is difficult in natural environments, as the number of possible interactions is high. An effective method to narrow down the list of potentially interacting microbes is via network analysis. Microbes that interact are likely to cooccur in the same habitat which in aquatic systems, unlike soils (Szoboszlay and Tebbe, 2021), can be spatially separated. Network analyses of communities can identify potential interacting partners and important metabolites (Mondav *et al*., 2020; Zamkovaya *et al*., 2020). Additionally, metabolic modelling and co-cultures can test hypotheses built by network analyses and vice-versa.

In this study we utilized a microbial community survey from a two-week depth discrete timeseries taken from Lake Lomtjärnan, Sweden. Lake Lomtjärnan is a small boreal lake with high methane concentrations in the water column. Because of this we hypothesized that a taxonomic-based network would reveal microbial consortia involved in C1 cycling. Furthermore, we could identify microbial phylotypes not previously documented as being involved in C1 cycling.

## Results and Discussion

### General observations

Our analysis of thirty-five 16S rRNA gene amplicon libraries yielded 123 840 OTUs. Of those, only 13 997 contained more than 2 sequences. Finally, the top 15 OTUs represented in average 51.25% of the relative abundance of the samples (Fig 2). While the surface and bottom of the water column had quite different dominant OTUs, the dominant prokaryotes shifted gradually along the water column. Only the upper water column showed changes across time supporting that the bottom waters were a more stable environment during the sampling campaign.

**Figure 2.**
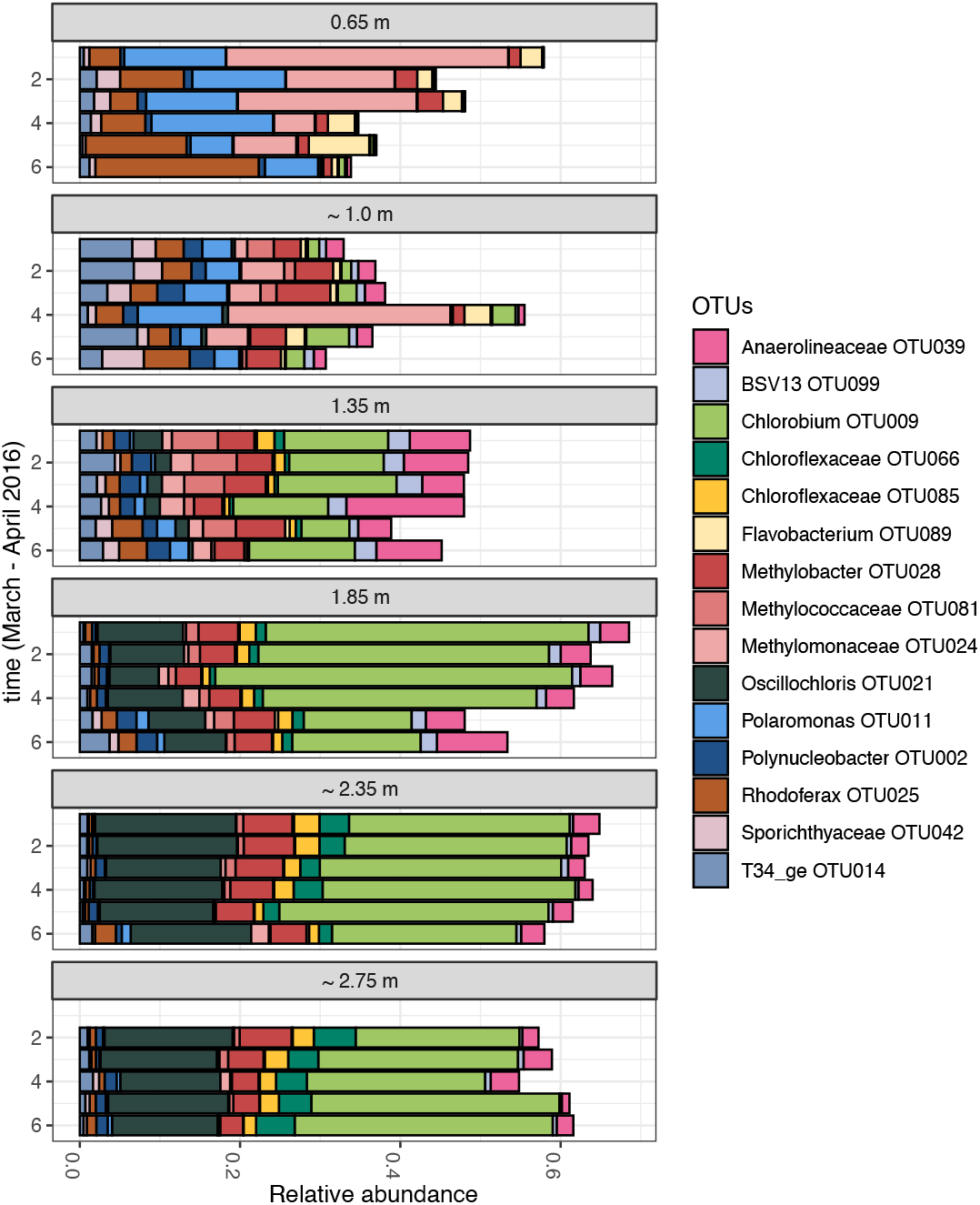
Fifteen most abundant OTUs from Lake Lomtjärnen time-depth series. OTUs are 95% identity clusters of V3-4 SSU rRNA gene region PCR amplicons taken from six depths from directly beneath the ice to the bottom of the water column. Six time points within 2 weeks (in average every other day) were collected during winter while the lake was ice covered.

### Lake-surface microbial consortia

Under ice-cover, aerobic β-proteobacterial *Rhodoferax* OTU025, *Polaromonas* OTU011, and γ-proteobacterial *Methylococcaceae* OTU024 (Fig 2, grey circle Fig 3) dominated Lake Lomtjärnen surface waters. All three potentially contribute directly to C1 cycling and are also predicted to fix N_2_ (Hanson *et al*., 2012; Bowman, 2016; Baker *et al*., 2017). *Rhodoferax* and *Polaromonas* are mixotrophs (Sizova and Panikov, 2007; Kato and Ohkuma, 2021; Taubert *et al*., 2022), being able to both fix inorganic and use organic C for energy and biomass. While *Methylococcaceae* are mostly reliant on CH_4_ (for both carbon and energy), some genera can supplement carbon acquisition via the reductive pentose phosphate cycle (CBB fixation) (Bowman, 2016). CH_4_ used by *Methylococcaceae* could be sediment derived or could come from nearby *Paludibacter* and *Clostridium* that use FeFe or VFe-type nitrogenases that have been shown to produce CH_4_ (Fixen *et al*., 2016; McRose *et al*., 2017; Addo and Dos Santos, 2020). The fact that the *Rhodoferax* phylotype was correlatively linked to *Ca*. Nomurabacteria is intriguing, given how little is known about the candidate genus. Comparing growth of *Ca*. Nomurabacteria on exudates from *Rhodoferax* compared to co-culturing could reveal if metabolite supply e.g. organic carbon, compared to syntrophic reliance e.g. direct electron transfer (Zhao *et al*., 2020) are occurring.

**Figure 3.**
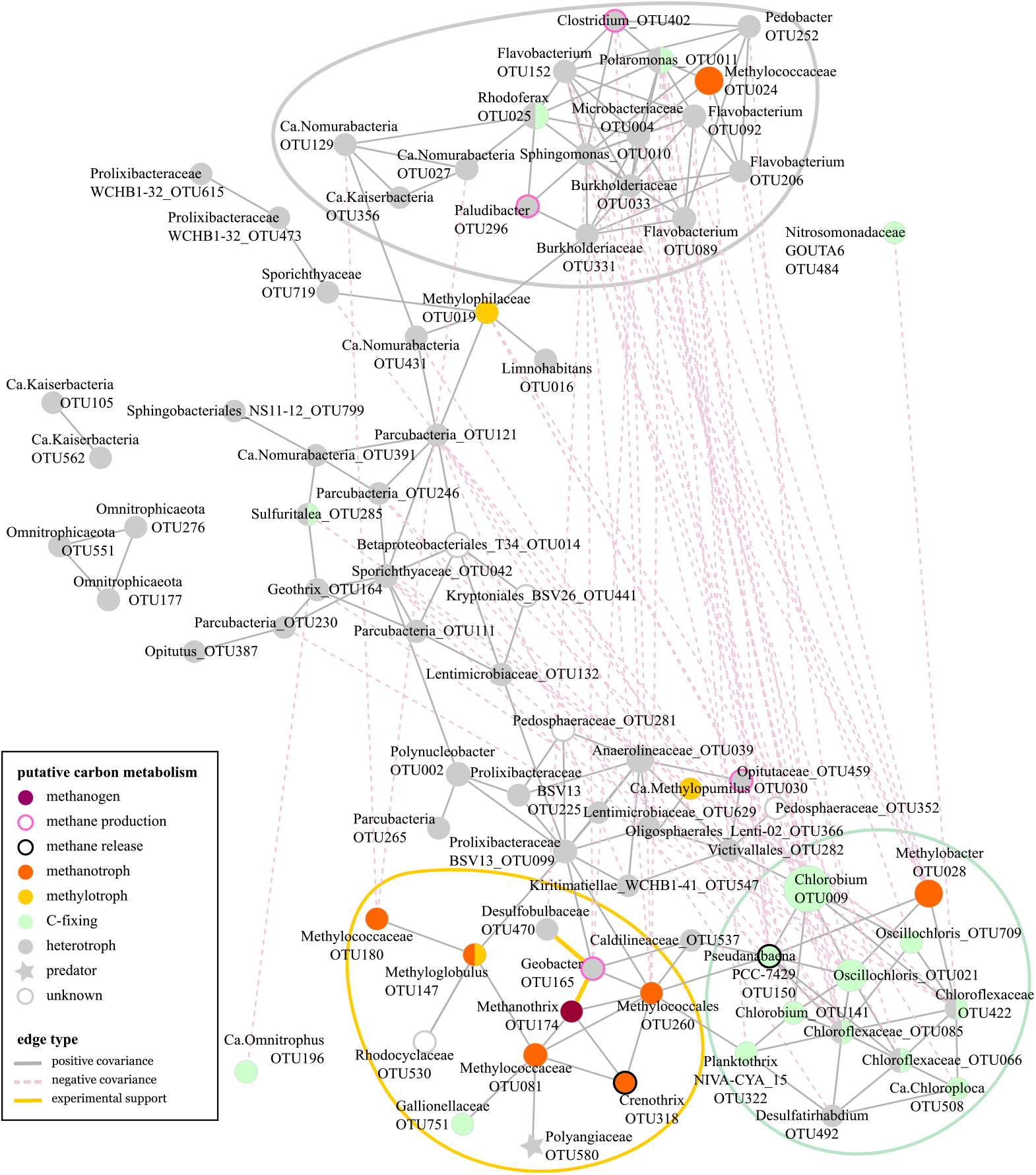
Lake time-depth-series network. Filled circles are OTUs coloured by putative carbon metabolism (SI_Table2 DOI: 10.6084/m9.figshare.21388026). Grey solid lines show positive correlations, grey dotted lines show negative correlations, and yellow lines designate experimentally supported interactions. Green, golden, and grey large circles designate potential consortia. Networks were calculated using 95% identity OTU abundance tables and at least two out of three consensus covariance method with SPARCC, SPIEC-EASI, and Pearsons correlations.

Most phylotypes in the grey circled cluster belong to taxa with aerobic to facultative anaerobic members and psychrotolerant or psychrophilic taxa (Finneran, 2003; Bernardet and Bowman, 2006; Darcy *et al*., 2011; Michaud *et al*., 2012; Viana *et al*., 2018). This cluster was associated with surface water which makes sense, as during winter the surface water, which is directly beneath the ice, is the coldest.

### Lake-bottom microbial consortia

*Chlorobium* OTU009, *Oscillochlorus* OTU021, and a *Methylobacter* OTU028 (Fig 2) dominated the deeper layers of the lake water. These three phylotypes are all obligate C1 cyclers. Network modules of deeper water microbes (green and gold circles Fig 5) consisted mostly of obligate or facultative C1 cyclers. *Planktothrix* OTU322 is a typical aerobic summer surface water inhabitant but as seen here it can ‘over-winter’ in deeper water and competes well in low-oxygen low-light conditions (Poulíčková *et al*., 2004; Ruuhijärvi *et al*., 2020). Another carbon fixing phylotype was *Pseudanabaeana* OTU150 which can release CH_4_ during photosynthesis and via Fenton-driven reactions (Cuellar-Bermudez *et al*., 2019). Fenton reactions can occur as intra or extracellular reactions whereby methylated compounds release CH_4_ in the presence of reactive oxygen species and a reduced transition metal, especially iron (Fe_2+_). Fenton reaction methane release is associated with organisms that can grow quickly and are heme prototrophs, especially those capable of oxygenic photosynthesis.

Active photosynthesising organisms, including *Cyanobacteria*, produce and export ROS and generate Fe_2+_ via both active heme-enzymes and during the production of the light harvesting photosystem. All that is needed for Fenton-driven CH_4_ release is the presence of methylated compounds e.g. DMSO, to explain the as yet unaccounted for component of (cyano/bacterial) aquatic methane (Bižić *et al*., 2020; Hilt *et al*., 2022). It is possible that all cellular lifeforms release CH_4_ especially during phases of rapid growth (Ernst *et al*., 2022) but in this study only bacteria with documented Fenton-driven release (Fig 3) are annotated. Although it cannot be ruled out that the *Methylobacter* phylotype is dependent on CH_4_ rising from methanogens in the sediments, links in the network analysis support partial reliance on bacterial CH_4._ Therefore, multiple lines of evidence point to adjacent phototrophs (*Chlorobium, Oscillochloris*, and *Chloroflexaceae*) as candidate CH_4_ cycling consortia clades worth further investigation.

In the golden circle the majority of C1 cyclers are methane cyclers with only *Gallionellaceae* annotated as a carbon fixer (Bethencourt *et al*., 2020). This methane cycling module also contains iron (*Geobacter, Crenothrix*, and *Gallionellaceae*) and sulfur cycling phylotypes (*Geobacter, Desulfobulbaceae* and *Gallionellaceae*) (Röling, 2014; Bethencourt *et al*., 2020). Both methanogens and methanotrophs have high iron and sulfur demand for cofactor production (Johnson *et al*., 2021). Of note is that the *Methanothrix* to *Geobacter* and *Geobacter* to *Desulfobulbaceae* network relationships are experimentally supported, where electron transfer between and from is the basis of the relationships (Holmes *et al*., 2004, 2017; Li *et al*., 2016; Liu *et al*., 2019; Song *et al*., 2020). The microbes in this module range from aerobic through to anaerobic and while the water column had some oxygen (Garcia *et al*., 2019) it is likely that the aerobic methanotrophs in bottom waters were assisted both by phototroph oxygen generation (van Grinsven *et al*., 2021) as well as CH_4_ release (Bižić *et al*., 2020). The presence of an archaeal methanogen in a potentially oxic environment might seem surprising, but *Methanothrix* (synonym *Methanosaeta* (Boone, 1991)) belongs to one of several methanogenic clades with oxygen tolerance (Jetten *et al*., 1989; Angle *et al*., 2017). The lack of genomic information available for many bacteria meant not all phylotypes could be assigned taxonomy at genus level. For this reason, predicting the metabolism of the *Rhodocyclaceae* phylotype was not possible as this family is particularly meta-bolically versatile. *Polyangiaceae* is a bacterial predator (Garcia and Müller, 2014), however, it is unlikely that its prey is the *Methylococcaceae* phylotype it covaries with, as predator-prey relationships result in negative covariance. It is unclear who is the *Polyangiaceae*s prey is as it has a broad host range (Petters *et al*., 2021), and a negative covariance with another OTU was not evident in the network. The *Polyangiacea*s role in this module may therefore be related to either direct production of nutrients via breakdown of complex molecules, or indirectly via release of nutrients from prey species. Of particular interest for future culture, co-culture/prey-baiting, and culture-independent studies are the three families: *Rhodocyclaceae, Gallionellaceae*, and *Polyangiaceae*.

Metabolic connections that likely tie these two modules together are: carbon fixation by the autotrophic and mixotrophic bacteria. These phylotypes also oxidise sulfur species to sulphate which could be consumed by the sulphate reducers. Methane is generated by methanogens and produced and released by phototrophs and consumed by the methanotrophs. Except for *Polyangiaceae* and *Pseudoanabaeana* the taxa in the green and gold circles are normally associated with low oxygen environments (Hanert, 2006; Pedersen, 2011; Garcia and Müller, 2014) supporting some role for habitat preference in the formation of these modules.

### Other water column microbes

*Polynucleobacter* OTU002 was highly abundant in lake water (Fig 2) showing no preference for depth. A large proportion of heterotrophs, especially those not associated with either the surface or bottom modules (Fig 3), were small-celled streamlined bacterial phyla *Patescibacteria* and *Omnitrophota*. Their specific contribution to carbon cycling is unspecified beyond most being heterotrophs. Given their position, ubiquity, and network position they likely facilitate carbon transfer between the surface and bottom waters. One *Omnitrophus* OTU196 could putatively fix CO2 (Kolinko *et al*., 2016) and was negatively corelated to the Geothrix OTU164. The methylotrophs *Ca*. Methylopumilus and *Methylophilaceae* found adjacent to the highlighted modules may play a critical role in reducing Fenton-driven methane release by mopping up methylated compounds.

### Taxonomic assignment of putative C1 cycling metabolism

Carbon fixation was once thought to be a purely phytoplankton (Cyanobacteria and eukaryotes) process achieved by coupling light-harvesting to Rubisco and the reductive pentose phosphate cycle. Over the last decade there have been some surprises in the taxonomic association of Rubisco, e.g., absence in *Melainabacteria* (Di Rienzi *et al*., 2013), presence in multiple Archaeal lineages, and non-fixation linked presence in DPANN archaea and *Patescibacteria* (Jaffe *et al*., 2019); But it is probable the metabolic potential of fixing carbon via Rubisco-CBB are now fully documented at least at the phylum level. However, at least six other well defined CO_2_ fixation pathways are known and likely many more deviations from canonical pathways will be elucidated as enzyme substitutions and pathway intermixing are discovered.

Determining that an organism is capable of producing or releasing methane is also complicated. There are many chemical reactions, enzymatic pathways, and taxonomic lineages involved (Liu *et al*., 2021)(Table 1). Here we used the term ‘methanogenesis’ to refer exclusively to methane generated via methyl-coenzyme M reductase (mcr) by archaeal methanogens and methanotrophs (Kevorkian *et al*., 2021). ‘Methane production’ here referred to the enzymatic generation of CH_4_ via FeFe or VFe type nitrogenases (Seefeldt *et al*., 2013; Zheng *et al*., 2018), and de-methylation of methylated compounds such as dimethyl sulfoxide (DMSO) and methyl phosphonates (Kamat *et al*., 2011; Klintzsch *et al*., 2019). The third category here was ‘methane release’ which covers the non-enzymatic generation of methane through Fenton-driven reactions and the breakdown of organic compounds in sunlight.

Methane is consumed aerobically by a few groups of bacteria. Two clades of archaea and one genus of bacteria oxidise it anaerobically (Welte *et al*., 2016). Community surveys of soluble and membrane bound (particulate) methane monooxygenase genes can identify bacteria capable of methane oxidation (bacterial methanotrophs) (Welte *et al*., 2016) while methyl-coenzyme M reductase surveys can identify archaeal methane oxidisers. However, their paraphyly, phylogenetic relatedness to archaeal methanogens, and the homology and reversibility of methanogenesis complicates identification of new archaeal methanotrophs (Bertram *et al*., 2013; Chadwick *et al*., 2022; Kevorkian *et al*., 2022).

## Conclusions

Contrary to historic understanding of carbon fixation in lakes, the surface consortia of an ice-covered lake consisted mostly of heterotrophs, and most of the photoautotrophs were in deeper water. The carbon fixing consortia was associated with another deep-water module mostly involved in CH4 cycling. Organic carbon produced by these consortia might transfer up the water column via streamlined bacteria and convection (Zdorovennova *et al*., 2021) to the surface waters feeding the cluster of heterotrophs found there (Fig 4). The most probable explanations for the negative correlation between the grey and gold-green circled modules is spatial separation controlled by water temperature, oxygenation, and lack of metabolic connection. However, with our approach, it is not possible to tell to what degree the co-occurrence of different phylotypes is due to shared environmental preference or by interactions.

**Figure 4.**
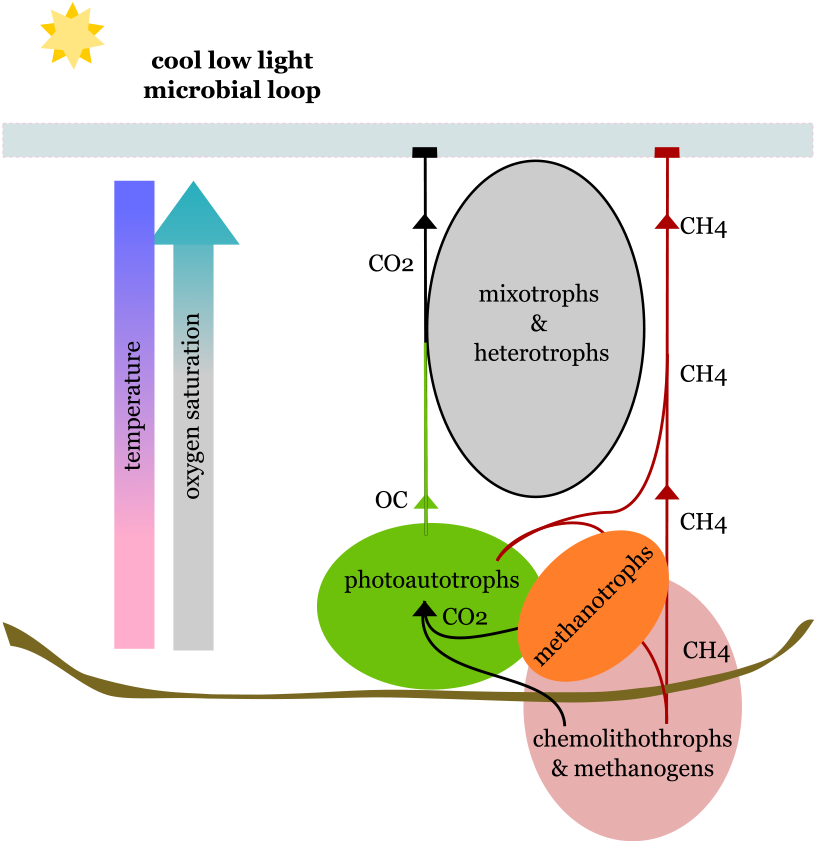
Summary diagram of winter microbial loop in Lake Lomtjärnen. the results from our study where we show that during ice-covered seasons carbon flows from the bottom of the water column to the surface.

Over the last two decades there has been a dramatic increase in the catalogued number of microbes and metabolic pathways capable of generating, releasing, or consuming C1 molecules (Alfreider *et al*., 2017; Steiger *et al*., 2017; Berghuis *et al*., 2019; Huang *et al*., 2019; Bižić *et al*., 2020; Kurth *et al*., 2020; Liu *et al*., 2021; Taubert *et al*., 2022). Such increases are likely to continue as we explore deeper genomic and metabolic space. Here, we identify six lineages (*Ca*. Nomurabacteria, *Chlorobium, Oscillochloris, Chloroflexaceae, Rhodocyclaceae, Gallionellaceae*, and *Polyangiaceae*) worth investigating for their roles in C1 cycling. A generalised call for greater focus on small-celled streamlined microbes in aquatic environments is also warranted. Follow up studies using a combination of meta’omics would clarify whether autotrophic or heterotrophic pathways are being utilised *in-situ*. Deliberate co-culture or reduced community culture from winter waters in cool low-light conditions is also worthwhile. In line with other studies, we found a distinct community of winter dominant microbes (Grzymski *et al*., 2012; Vigneron *et al*., 2019). Despite the difficulties of sampling in winter (Block *et al*., 2019) a more equal percentage of studies of inland water-bodies need to focus on under ice microbiomes if we are to truly understand aquatic C1 cycling.

### Data availability and source code

All scripts used to calculate network correlations are available at: https://github.com/rmondav/C1_cycling_under_ice, v1.0.0 recorded at https://doi.org/10.5281/zenodo.5531947. Time-depth-series amplicons are available under BioProject PRJEB27633 (Garcia *et al*., 2019). SI Tables 1 is available at https://doi.org/10.6084/m9.figshare.21388005 and SI Table 2 at https://doi.org/10.6084/m9.figshare.21388026.

## Acknowledgments

SLG and SP were each supported by their own SciLifeLab fellowships. RM was supported by a Swartz Scholarship.

Computations were enabled by resources in projects 2020-5-529 & 2020-15-261 and data storage projects 2020-6-164, 2020-16-196, & 2021-23-646 provided by the Swedish National Infrastructure for Computing (SNIC) at UPPMAX, which is partially funded by the Swedish Research Council through grant agreement no. 2018-05973. SLG obtained financial resources while SP and RM obtained computational resources. GM, RM, and SLG wrote and/or implemented scripts for analysis and visualization of results. RM and SP drafted the manuscript and all authors contributed to discussions, editing, and approval of final manuscript. RM curated scripts. SP and SLG supervised.

