## Supplementary material for "Network analysis uncovers associations in the turnover of C1 molecules in a winter lake": SI_Materials_and_Methods

[*] Current address: Swedish Nuclear Fuel and Waste Management Co.

Experimental Procedures ........................................ 2
References ........................................ 4

### Experimental Procedures

*Lake water collection and media preparation*

Samples were collected from Lake Lomtjärnan located in the central west (Jämtland) of Sweden (Fig S1). The surface area of the lake is about 1 ha, and the maximum depth is 3.5 m. The lake is located on a mire surrounded by a coniferous forest. Samples for the time-depth-series were collected at six time points in winter from the last week of March to the first week of April, 2016 and are published in Garcia *et al*, 2019. The lake was ice covered throughout the sampling campaign. At each sampling occasion, samples at six different depths (0.65 m, 1.0 m, 1.35 m, 1.85 m, 2.35 m, 2.75 m) were taken, totalling 35 samples. The deepest depth was not taken on the first sampling because the max depth was around 2.35 m at that location at that time (Garcia *et al.*, 2019). Water for the time-depth-series was collected using a depth-discrete Limnos tube-sampler (Limnos, Poland) and the water was subsequently filtered through a Sterivex filter (0.22 µm) and the filters were stored immediately in liquid nitrogen for later DNA extraction.

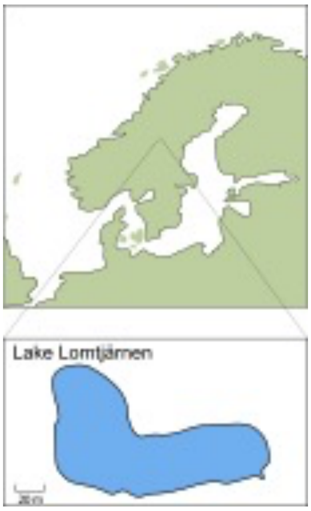


Figure S1. Map of nordic region showing location of Lake Lomtjärnen

*16S rRNA gene amplicon preparation and sequencing*

Total nucleic acids were extracted using a phenol-chloroform method (Griffiths *et al.*, 2000) with modifications (Taipale *et al.*, 2009). Library preparation for 16S rRNA gene analysis was done following a two-step Polymerase Chain Reaction (PCR) protocol performed in triplicate with primers 341F (3'-CCTACGGGNGGCWGCAG-5') (Herlemann *et al.*, 2011) and 805NR (3'-GACTACNVGGGTATCTAA-5') (Mondav *et al.*, 2020). All PCRs were conducted in 20 µl volume using 0.02 U/µl Phusion high fidelity DNA polymerase, 1X Q5 reaction buffer (NEB, UK), 0.25 µM primers and 200 µM dNTP mix and 1 µl template. The thermal cycler program consisted of an initial 98°C denaturation step for 10 min followed by 20 cycles of denaturation at 98°C for 10 seconds, annealing at 48°C for 30 seconds, and polymerisation at 72°C for 30 seconds with a final elongation step at 72°C for 2 minutes. The triplicate PCR reactions were pooled then purified with magnetic beads (Sera-Mag™ Select, GE Healthcare, Chicago, United States of America), and 2 µl of the purified product used as template for a second stage PCR to add indexed primers. The second thermal program consisted of an initial 98°C denaturation step for 30 seconds, then 15 cycles of 98°C for 10 seconds, 66°C for 30 seconds, and 72°C for 30 seconds and a final elongation at 72°C for 2 minutes. Following index addition PCR products were again purified with magnetic beads and quantified with Qubit™ using the Qubit™ dsDNA HS Assay Kit (Invitrogen™). Finally, 15.6 µg of each indexed and purified PCR product were pooled before submission of the sample to the Science for Life Laboratory SNP/SEQ sequencing facility hosted by Uppsala University (Uppsala, Sweden). Sequencing was done using Illumina Miseq in paired-end mode with 300bp and v3 chemistry.

*16S rRNA gene amplicon processing and bioinformatics*
Sequence processing was performed with Mothur 1.41.0 following the MiSeq SOP (Kozich *et al.*, 2013), with the exception that clustering to operational taxonomic units (OTUs) was done using VSEARCH (Rognes *et al.*, 2016) as implemented in Mothur (Schloss *et al.*, 2009). Taxonomy was assigned against the Silva 132 database (Quast *et al.*, 2013), sequences were then subsampled once using Qiimes (Caporaso *et al.*, 2010) single rarefaction to an even depth of 2000 reads per sample. The normalized OTU table was used for compositional and comparative analyses. Column graphs of the ten most abundant phylotypes were made for compositional analyses. OTU tables were processed and visualised in R 4.0.3 (R Core Development Team, 2015) using phyloseq 1.34.0 (McMurdie and Holmes, 2013) and ggplot2 (Ginestet, 2011). Prior to co-occurrence analysis for network visualisation, OTUs present in less than 50% of the lake water samples, or with a relative abundance always below 1 % were removed. This was done to reduce sparcity to below 50%. A network ensemble approach was used where any correlation (edge) between OTUs (node) that was found in at least two of the three following methods was included in the final visualisation: SPIEC-EASI (Kurtz *et al.*, 2015), SparcCC (Friedman and Alm, 2012), and Pearsons correlation. All co-occurrence methods were implemented in R 4.0.3 (R Core Development Team, 2015) using spiec-easi 1.1.1 for SPIEC-EASI and SparcCC, and the psych 2.1.3 package for Pearsons correlation. Networks were visualised in Cytoscape. Inkscape was used to manually add color-coded hemispheres to OTU nodes and literature-based notations and colouring of lines between nodes (edges). Labels were moved to avoid overlap. Metabolic predictions based on taxonomy were gained by literature review and keyword search of PubMed, Mendelay, and Google-Scholar databases. Taxonomic information on the *nifD* components of nitrogenases TIGR01861 (Fe-Fe), TIGR01860 (V-Fe), and IPR010143 (Mo-Fe) that predict methane production potential was obtained from UniProt-Interpro (Haft D.H. *et al.*, 2003; Magrane and Consortium, 2011; Punta *et al.*, 2012).
